# ConformFlow: scalable normalizing flow for protein conformational ensemble generation

**DOI:** 10.64898/2026.06.12.731927

**Authors:** Yikai Liu, Guang Lin, Ming Chen

## Abstract

Molecular dynamics (MD) simulations remain the standard tool for characterizing protein conformational landscapes, but their high computational cost limits large-scale and long-timescale applications. Recent generative models, especially diffusion-based approaches, provide promising alternatives by learning equilibrium conformational distributions across diverse protein systems. We present **ConformFlow**, the first scalable normalizing-flow framework for sequence-conditioned protein conformational ensemble generation. ConformFlow combines a continuous backbone latent representation with a RealNVP-style flow parameterized by sequence-aware Transformer coupling networks, enabling exact likelihood training, single-step sampling, and plug-and-play conditioning on flexible geometric constraints. Across diverse protein systems, ConformFlow generates ensembles that agree well with reference MD simulations, generalizes to proteins beyond its training data, and achieves substantially faster sampling than diffusion-based baselines. These results establish ConformFlow as an efficient and controllable alternative for protein conformational ensemble generation.

**Code:** https://github.com/Harrydirk41/ConformFlow.git

## 1 Introduction

Proteins are essential molecular components of living systems, carrying out diverse functions such as catalysis, signaling, and transport. These functions are governed not only by static structures, but also by conformational flexibility and dynamic transitions. Characterizing protein conformational ensembles is therefore critical for understanding the molecular mechanisms underlying protein function Whisstock & Lesk (2003). Molecular dynamics (MD) simulations provide a principled approach for studying biomolecular processes such as protein folding and unfolding Shaw et al. (2010); Robustelli et al. (2018). However, MD remains computationally demanding, making exhaustive simulation of long-timescale protein dynamics computationally prohibitive.

Recent progress in deep generative modeling has opened new opportunities for protein conformational modeling. A growing body of work has developed generative approaches for sampling equilibrium ensembles Lu et al. (2023); Lewis et al. (2025); Jing et al. (2024a); Noé et al. (2019); Zheng et al. (2024); Wayment-Steele et al. (2024); Jing et al. (2024b); Raja et al. (2025); Lelièvre et al. (2023); Du et al. (2024); Fu et al. (2022); Arts et al. (2023); Liu et al. (2025b). These methods offer a promising route to studying protein–protein interactions, allosteric regulation, biomolecular condensation, and conformational heterogeneity Brandsdal & Smalås (2000); Guo & Zhou (2016).

Most recent scalable approaches are based on diffusion models, which are flexible, expressive, and have shown promising generalization across diverse protein systems. However, diffusion models require iterative denoising at inference time, making sampling less efficient. Moreover, because exact likelihood evaluation is generally unavailable for diffusion models, incorporating likelihood-based physical constraints or posterior guidance is less direct.

Motivated by these opportunities and limitations, we introduce **ConformFlow**, a Normalizing-Flow framework for protein Conformational ensemble generation. To our knowledge, ConformFlow is the first scalable normalizing-flow framework for sequence-conditioned protein conformational ensemble generation. It combines a learned structural latent representation with a scalable RealNVP-style generative model parameterized by masked Transformer coupling networks. The model is trained on large-scale single-structure datasets together with equilibrium MD trajectories, allowing it to learn transferable distributions over protein conformational space. Unlike diffusion-based models, ConformFlow provides single-step sampling and exact likelihood evaluation. These properties make it computationally efficient and naturally compatible with inference-time guidance under flexible geometric constraints.

Our contributions are fivefold:

1. **A scalable architecture for sequence-conditioned normalizing flow**. We introduce a latent-space RealNVP architecture with residue-masked affine coupling layers parameterized by bidirectional Transformer as backbones, enabling expressive sequence-conditioned density modeling while preserving parallel sampling and exact likelihood evaluation.
2. **Large-scale transferable training**. We train ConformFlow on hundreds of thousands of protein sequences and millions of conformations from single-structure databases and equilibrium MD trajectories, enabling ensemble generation across diverse protein families.
3. **Fast and controllable inference**. ConformFlow enables one-step generation with exact likelihood evaluation, yielding substantially faster sampling than iterative diffusion models and supporting plug-and-play guidance with flexible, user-specified geometric constraints.
4. **Scaling behavior of flow depth**. We show that ConformFlow benefits substantially from increasing the number of coupling blocks, improving ensemble fidelity on training proteins while also enhancing generalization to held-out protein systems.
5. **Ablation study of NF architecture choices**. We systematically ablate coordinate representation and coupling-mask design, showing that the learned latent representation and odd–even residue coupling are both important for accurate and transferable sequence-conditioned protein ensemble generation.

## 2 Background

### 2.1 Normalizing Flows and Realnvp

Normalizing flows are likelihood-based generative models for continuous random variables. Let *x* ∈ℝ^*D*^ denote a data sample drawn from an unknown data distribution *p*_data_. A normalizing flow defines an invertible and differentiable transformation

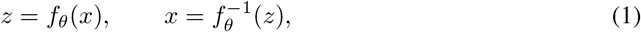

where *z* is mapped to a simple base distribution *p*_0_(*z*), typically a standard Gaussian *N* (0, *I*). By the change-of-variables formula, the model density is

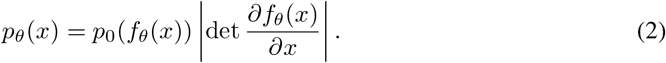

Maximum-likelihood training therefore minimizes the negative log-likelihood

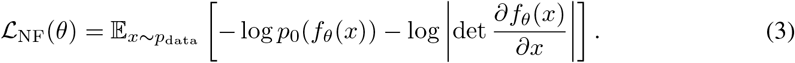

When *p*_0_(*z*) = *N* (0, *I*), this objective can be written, up to an additive constant, as

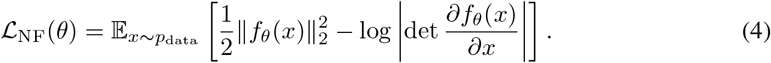

The first term encourages data samples to be mapped to high-probability regions of the base distribution, while the second term accounts for the local volume change induced by the transformation. After training, generation is performed exactly by sampling *z ∼ p*_0_(*z*) and applying the inverse map 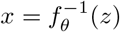.

A key challenge in normalizing flows is designing transformations that are both expressive and have tractable Jacobian determinants. RealNVP Dinh et al. (2016) addresses this using affine coupling layers. Given an input *x*, split it into two parts *x*_*a*_ and *x*_*b*_. A coupling layer keeps one part unchanged and affinely transforms the other:

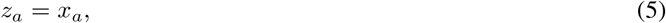

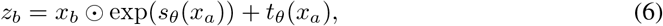

where *s*_*θ*_ and *t*_*θ*_ are neural networks producing scale and translation parameters. The inverse transformation is analytic:

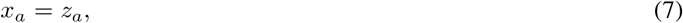

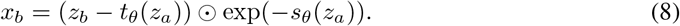

Because the Jacobian of this transformation is triangular, its log-determinant is cheap to compute:

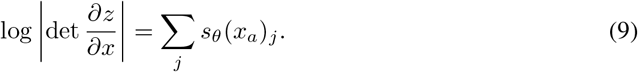

Stacking multiple coupling layers with alternating masks or permutations allows all dimensions to be transformed while preserving exact likelihood evaluation and exact sampling.

Historically, normalizing flows have faced a tradeoff between expressivity and tractable Jacobians. Coupling-based models such as RealNVP are efficient, but each layer only transforms part of the input, which can limit expressivity. Autoregressive flows Papamakarios et al. (2017) improve expressivity by allowing each dimension or block to depend on previous ones while preserving a triangular Jacobian. Recent models such as TARFLOW Zhai et al. (2024) scale this idea with causal Transformer blocks. However, inverse sampling becomes sequential along the autoregressive order, making generation slower than parallel coupling-based flows.

### 2.2 Deep Generative Modeling for Protein Conformational Ensembles

Deep generative models have recently been applied to protein conformational ensemble generation by learning sequence-conditioned distributions over structures from structural databases or equilibrium MD trajectories. Early likelihood-based approaches used normalizing flows to model equilibrium molecular distributions Kohler et al. (2023). These methods were mainly demonstrated on individual molecular systems and did not establish scalability or transferability across diverse protein sequences. Most recent scalable ensemble generators instead adopt diffusion- or score-based formulations Lu et al. (2023); Lewis et al. (2025); Jing et al. (2024a); Noé et al. (2019); Zheng et al. (2024); Wayment-Steele et al. (2024). These models have shown strong performance by learning transferable conformational distributions across broad protein families. At the same time, the lack of exact likelihood has motivated renewed interest in modern normalizing-flow architectures for transferable learning across molecular systems Tan et al. (2026); Klein & Noé(2024). However, existing NF-based approaches remain primarily focused on peptides and small proteins, and have not yet demonstrated large-scale sequence-conditioned protein ensemble generation.

### 2.3 Protein Conformational Representations

Recent advances have shown that protein conformations can be effectively represented through learned structural representations, ranging from residue-level discrete tokenizers to continuous coordinate encoders. For example, ESM3 introduces a structure tokenizer that maps a protein conformation to a discrete token sequence. In parallel, SimpleFold uses continuous atom-level coordinate encoders within a flow-matching framework Wang et al. (2025). These representations provide compact alternatives to directly modeling raw three-dimensional coordinates and can serve as intermediate spaces for generative modeling. In ConformFlow, we adopt a continuous structural latent space, allowing the NF model to perform exact likelihood-based density estimation in a latent representation while retaining a decoder back to protein coordinates.

## 3 Method

In this section, we provide a high-level overview of the ConformFlow architecture. ConformFlow consists of two components. First, a structure autoencoder maps backbone Cartesian coordinates into a compact continuous latent representation. Second, a sequence-conditioned normalizing flow models the distribution over this latent space using RealNVP-style coupling layers parameterized by modern masked Transformer networks. An overview is shown in Fig. 1.

**Figure 1.**
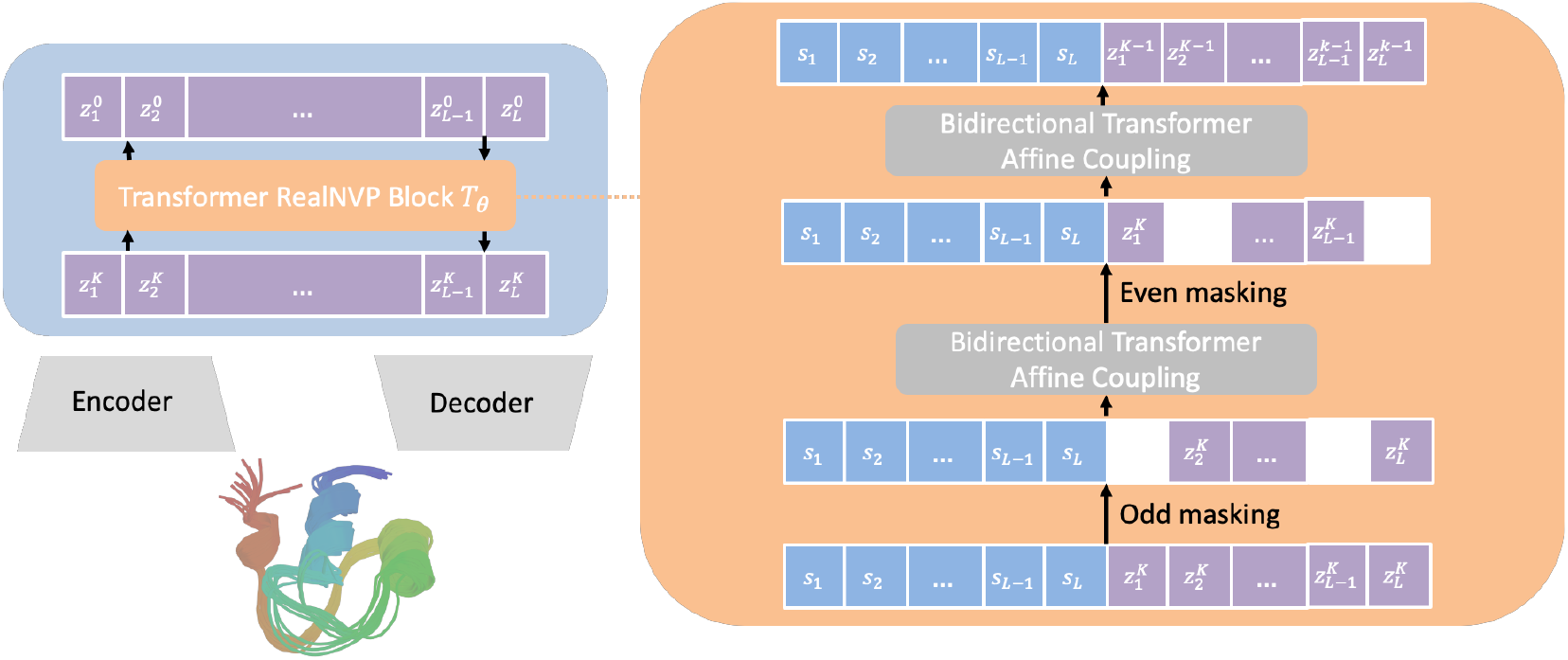
Overview of **ConformFlow.** ConformFlow (1) maps backbone Cartesian coordinates **x** ∈ℝ^*L×*3*×*3^, consisting of N, C*α*, and C atoms, into a compact continuous latent representation 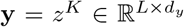 using an autoencoder model. (2) A sequence-conditioned RealNVP then learns the latent ensemble distribution *p*_*θ*_(**y** | *s*) through a stack of invertible affine coupling blocks. During sampling, Gaussian noise is transformed through the invertible flow into latent conformations, which are then decoded back to backbone coordinates. Each coupling block keeps part of the latent variables fixed under an odd–even masking scheme and updates the remaining variables using bidirectional transformer coupling networks conditioned on the amino-acid sequence *s*.

### 3.1 STRUCTURE AUTOENCODER

Let **x** ∈ℝ^*L×*3*×*3^ denote the backbone coordinates of a protein with *L* residues, where each residue contains the atoms {*N, C*_*α*_, *C*}. We encode each conformation into a continuous per-residue latent representation

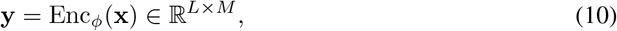

where *M* is the latent dimension per residue.

Our autoencoder is adapted from the ESM3 structure tokenizer. The original ESM3 tokenizer maps backbone coordinates to discrete structure tokens through a vector-quantized bottleneck. In contrast, we replace the vector quantizer with a continuous bottleneck. Specifically, the frozen ESM3 structure encoder produces per-residue features in ℝ^1280^, which are mapped by a trainable adapter into ℝ^*M*^. The decoder applies a second trainable adapter from ℝ^*M*^ back to ℝ^1280^, followed by the frozen ESM3 structure decoder:

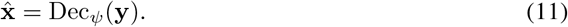

Thus, the pretrained geometric encoder and decoder are kept fixed, while only the lightweight bottleneck adapters are trained. This design preserves the geometric inductive biases of ESM3 while producing a continuous latent space suitable for likelihood-based modeling. Additionally, because the continuous latent representation is derived from the ESM3 structure tokenizer, it inherits invariance to global SE(3) transformations of the input structure.

We train the autoencoder to reconstruct backbone geometry using the first four geometric Stage-1 objectives of ESM3:

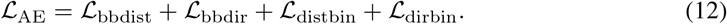

Here, ℒ_bbdist_ preserves pairwise backbone-atom distances, ℒ_bbdir_ preserves relative backbone directions, ℒ_distbin_ predicts discretized *C*_*β*_–*C*_*β*_ distance bins, and ℒ_dirbin_ predicts discretized directional bins. Together, these losses encourage the continuous latent representation to preserve both metric and orientational information needed for accurate backbone reconstruction.

### 3.2 Sequence-conditioned latent normalizing flow

Once the autoencoder is trained, we freeze it and learn a sequence-conditioned density over the continuous latent representation,

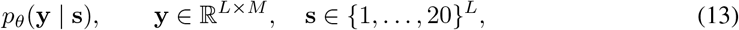

where *L* is the number of residues, *M* is the latent dimension per residue, and **s** is the amino-acid sequence. The flow maps a standard Gaussian base variable to a protein latent:

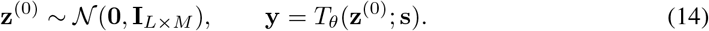

The transformation *T*_*θ*_ is a stack of *K* invertible blocks,

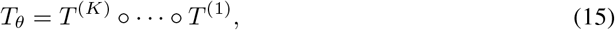

with intermediate states

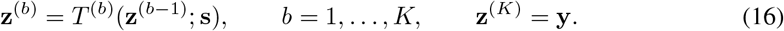

#### Residue-masked affine coupling

Each block *T* ^(*b*)^ contains two affine coupling layers. The first updates the odd-indexed residues while keeping the even-indexed residues fixed; the second updates the even-indexed residues while keeping the odd-indexed residues fixed:

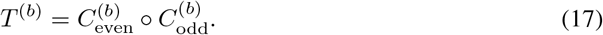

Thus, within each block, every residue latent vector is updated once. Stacking multiple blocks yields a deep sequence-conditioned bijection over the full protein latent representation.

For a generic coupling layer *C*_*m*_, let Ω_*m*_ ⊂ { 1, …, *L*} denote the active residues to be updated and let

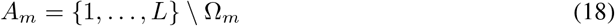

denote the frozen residues. Given the current state **z** ∈ℝ^*L×M*^, the coupling network observes only the frozen residue latents 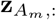, together with the full sequence embedding **e**(**s**). It predicts scale and translation fields for the active residues:

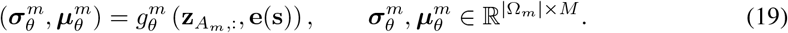

The active residue latents are then transformed by a per-residue affine map:

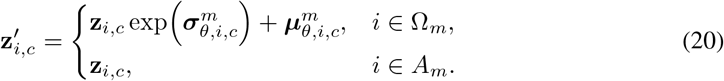

The conditioner 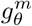 is implemented as a bidirectional masked Transformer coupling network. It attends bidirectionally across residue positions, using the full amino-acid sequence embedding and only the frozen residue latents as input. The active residue latents are masked from the conditioner, preserving the triangular Jacobian required for exact likelihood computation. The final output head predicts the scale and translation fields jointly.

#### Exact likelihood training

Because each coupling layer transforms only the active residues using parameters computed from the frozen residues, the Jacobian of Eq. equation 20 is block triangular. Its log-determinant is

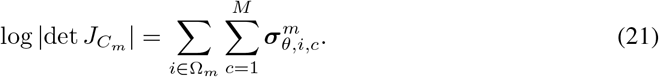

The full-flow log-determinant is the sum over all coupling layers and blocks.

We train the flow by exact maximum likelihood on latents by change of variables,

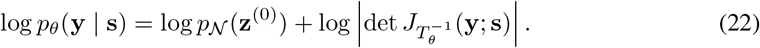

#### Sampling

To generate an ensemble for a sequence **s**, we sample

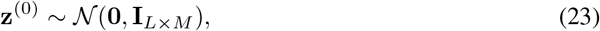

apply the forward flow

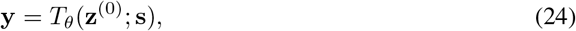

and decode the latent conformation to backbone coordinates:

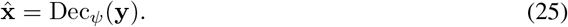

### 3.3 Plug-and-play conditional generation

ConformFlow supports inference-time control by combining the learned sequence-conditioned density with differentiable geometric constraints. Let *f* : ℝ^*L×*3*×*3^ →ℝ be a user-specified differentiable observable, such as radius of gyration or pairwise distance, and let *v* be a target value. We define a soft-constrained posterior over latent conformations:

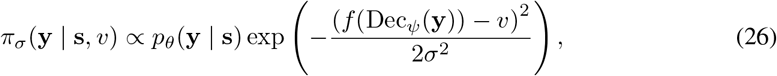

where *σ* controls the constraint strength.

We sample from this posterior using unadjusted Langevin dynamics. In latent space, the update is

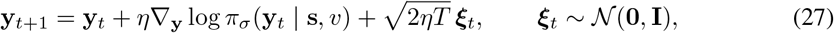

where *η* is the step size and *T* is an optional sampling temperature.

Alternatively, we can sample in the Gaussian base space by writing

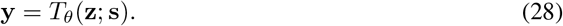

This defines the corresponding target

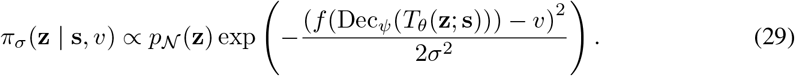

In both parameterizations, *f* only needs to be differentiable, so new geometric constraints can be introduced at inference time without modifying or retraining the model.

## 4 EXPERIMENT

We evaluate ConformFlow on protein conformational ensemble generation across diverse protein systems. Our results demonstrate that ConformFlow accurately recovers equilibrium distributions from reference MD simulations, generalizes to unseen proteins systems, and enables substantially faster sampling. We further show that the learned likelihood-based model supports plug-and-play conditional generation under differentiable geometric constraints without retraining. Finally, we demonstrate comprehensive ablation studies on scaling behaviors and key architecture design.

### 4.1 Experimental setup

#### Data

For training the continuous structure autoencoder, we use the Swiss-Prot subset of the AlphaFold Database, containing 542,378 protein predicted native structures. This dataset provides broad coverage of protein structures and is used to learn latent representations of geometry.

For training the sequence-conditioned normalizing flow, we combine single-structure data with equilibrium MD trajectories. The single-structure component uses the Swiss-Prot subset of the AlphaFold Database. The MD component combines mdCATH Mirarchi et al. (2024) with the BioEmu simulation corpus Lewis et al. (2025), including CATH1, CATH2, Octapeptides, and MEGAsim Charron et al. (2025); Sillitoe et al. (2021); Tsuboyama et al. (2023). ATLAS Vander Meersche et al. (2024) is held out and used only as test dataset.

#### Model details

ConformFlow is trained on precomputed continuous latents, with the structure autoencoder frozen during flow training. We use latent dimension *M* = 4 per residue and a 12-block RealNVP model with Transformer coupling networks. Additional training details are provided in Appendix B.1.

#### Baselines

Our main baseline is BioEmu Lewis et al. (2025), a recent state-of-the-art model for transferable protein conformational ensemble generation trained at comparable scale. Although several diffusion- and flow-matching-based methods have been proposed for ensemble generation, BioEmu is among the few models trained on large-scale, high-quality MD simulation data and is therefore the most direct comparison for our setting.

We also compare against normalizing-flow-based baselines in the ablation study. While prior NF-based molecular ensemble generators have demonstrated promising results, they have primarily focused on small molecules, peptides, or small protein systems. To evaluate the effect of NF architecture design under our large-scale training setting, we adapt PROSE Tan et al. (2026), a TARFlow-based architecture, and retain its major architectural components while training it on our dataset. This provides a controlled NF baseline for assessing the impact of the learned latent representation and coupling-mask design used in ConformFlow.

### 4.2 Results

#### Distributional similarity

We evaluate ensemble quality by comparing generated conformational distributions against reference MD simulations. We report Jensen–Shannon divergence (JSD) between reference and generated ensembles projected onto four collective variables: end-to-end distance (EED), radius of gyration (*R*_*g*_), RMSD to the native structure, and the top two TICA components computed from *C*_*α*_ pseudo-torsion angles Schultze & Grubmuuller (2021).

We evaluate three settings: CATH1, a seen-protein benchmark with 50 proteins; held-out octapeptides, an unseen sequence benchmark in a length-matched regime; and ATLAS, an unseen-protein benchmark outside the training corpus Vander Meersche et al. (2024). Table 1 summarizes the results. On CATH1, ConformFlow improves EED, RMSD, and 2D TICA while matching BioEmu on *R*_*g*_. On held-out octapeptides, ConformFlow achieves consistently lower JSD across all reported collective variables, demonstrating generalization to unseen sequences in the peptide regime. On ATLAS, ConformFlow also improves all metrics despite the dataset being fully held out from training, indicating transferability to external protein simulation data. Overall, these results show that ConformFlow captures both low-dimensional geometric observables and slow collective modes across seen, held-out, and external test settings. We provide visual free-energy comparisons on representative CATH1 systems in Appendix Figures 2 and 3. ConformFlow recovers the main low-free-energy regions observed in reference MD and closely tracks the MD landscapes.

**Table 1.** JS divergence against MD and wall-time on test proteins from each dataset.

| Quantity | CATH1 ($N = 50$) | | Octapeptide ($N = 50$) | | ATLAS ($N = 40$) | |
| --- | --- | --- | --- | --- | --- | --- |
|  | BioEmu | ConformFlow | BioEmu | ConformFlow | BioEmu | ConformFlow |
| End-to-end | 0.062 (0.006) | **0.043 (0.001)** | 0.039 (0.000) | **0.035 (0.000)** | 0.291 (0.005) | **0.255 (0.006)** |
| $R_g$ | **0.083 (0.007)** | 0.085 (0.004) | 0.038 (0.000) | **0.032 (0.000)** | 0.323 (0.019) | **0.305 (0.010)** |
| RMSD | 0.124 (0.012) | **0.058 (0.003)** | 0.038 (0.000) | **0.033 (0.000)** | 0.366 (0.032) | **0.330 (0.023)** |
| 2D TICA | 0.182 (0.011) | **0.132 (0.003)** | 0.336 (0.005) | **0.329 (0.004)** | 0.434 (0.001) | **0.403 (0.000)** |
| Time (s) | 451.9 | **10.7** | 71.1 | **4.7** | 2181.1 | **25.8** |

#### Plug-and-play conditional generation

One important advantage of diffusion-based generative models is their flexibility for inference-time guidance Ho & Salimans (2022); Chung et al. (2022). Prior work has shown that diffusion-based ensemble generators can be guided by physical forces Wang et al. (2024) and experimental constraints Liu et al. (2025a), enabling conditional generation without retraining. In this study, we evaluate whether ConformFlow can be controlled at inference time without retraining. We use RMSD to the native structure as a differentiable constraint observable and draw target values from the empirical MD RMSD distribution. We compare ULA guidance in the learned latent space **y** and in the Gaussian base space **z**, denoted ConformFlow (**y**-guided) and ConformFlow (**z**-guided), respectively. Implementation details are provided in Appendix B.2.

Table 2 reports results on ATLAS. Both guided samplers improve agreement with reference MD across all evaluated collective variables, showing that knowledge of low-dimensional features can enhance ensemble generation quality. The **z**-guided sampler performs best overall. Intuitively, applying Langevin updates in the gaussian base space avoids sampling on a rugged free energy surface and leads to smoother exploration. This is also a practical and unique advantage of ConformFlow over autoregressive flows: Gaussian-space guidance would require mapping each updated base-space sample back to the latent space at every step, which is sequential and expensive for autoregressive models. In contrast, ConformFlow’s RealNVP-style map is parallel, making **z**-space guidance practical for long protein sequences.

**Table 2.** Effect of plug-and-play RMSD guidance on ConformFlow ensembles, evaluated on ATLAS dataset. Guidance targets are drawn from the MD empirical RMSD distribution, so the RMSD JSD measures constraint satisfaction for the guided variants.

|  | ConformFlow | ConformFlow (y-guided) | ConformFlow (z-guided) |
| --- | --- | --- | --- |
| End-to-end | 0.255 (0.006) | 0.209 (0.008) | **0.195 (0.003)** |
| $R_g$ | 0.305 (0.010) | 0.177 (0.011) | **0.145 (0.008)** |
| RMSD | 0.330 (0.023) | 0.061 (0.009) | **0.043 (0.007)** |
| 2D TICA | 0.403 (0.000) | 0.348 (0.002) | **0.312 (0.004)** |

#### Scaling behavior

We study how ConformFlow performance changes as the number of affine coupling blocks increases. Table 3 reports JSD against reference MD for models with 4, 8, and 12 coupling blocks. Increasing flow depth consistently improves distributional similarity across datasets and collective variables. The gains are especially pronounced on held-out octapeptides and ATLAS, suggesting that deeper coupling stacks improve not only fitting of training-related protein ensembles, but also transfer to unseen protein systems.

**Table 3.**
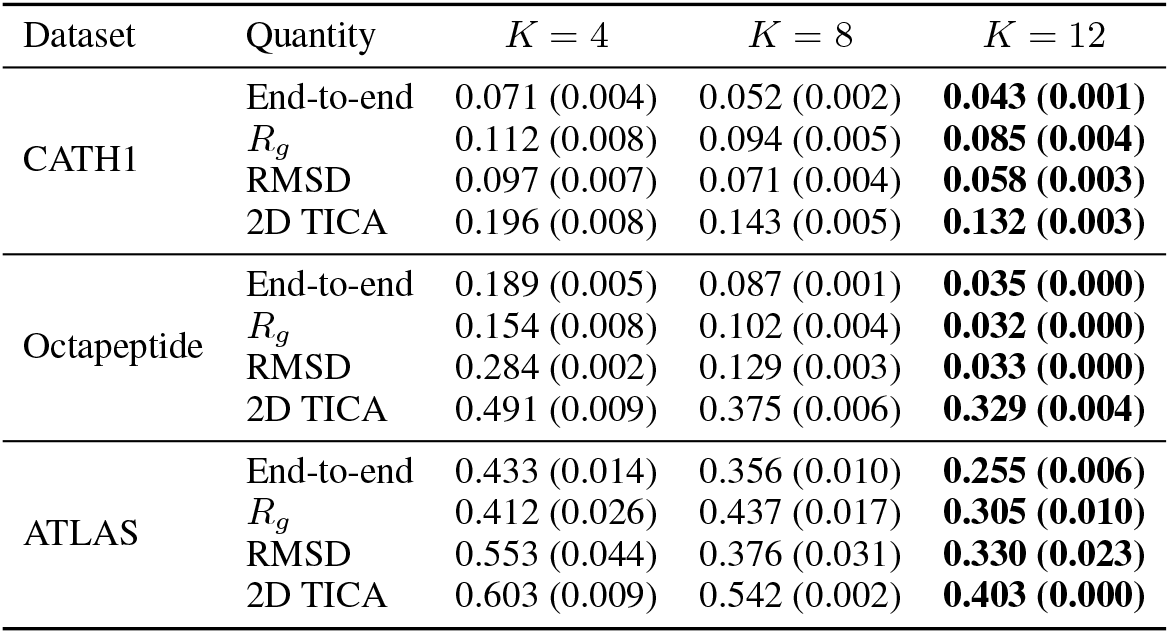
JS divergence against MD on test proteins from each dataset for different numbers of coupling blocks.

| Dataset | Quantity | $K = 4$ | $K = 8$ | $K = 12$ |
| --- | --- | --- | --- | --- |
| CATH1 | End-to-end | 0.071 (0.004) | 0.052 (0.002) | **0.043 (0.001)** |
| | $R_g$ | 0.112 (0.008) | 0.094 (0.005) | **0.085 (0.004)** |
|  | RMSD | 0.097 (0.007) | 0.071 (0.004) | **0.058 (0.003)** |
|  | 2D TICA | 0.196 (0.008) | 0.143 (0.005) | **0.132 (0.003)** |
| Octapeptide | End-to-end | 0.189 (0.005) | 0.087 (0.001) | **0.035 (0.000)** |
| | $R_g$ | 0.154 (0.008) | 0.102 (0.004) | **0.032 (0.000)** |
|  | RMSD | 0.284 (0.002) | 0.129 (0.003) | **0.033 (0.000)** |
|  | 2D TICA | 0.491 (0.009) | 0.375 (0.006) | **0.329 (0.004)** |
| ATLAS | End-to-end | 0.433 (0.014) | 0.356 (0.010) | **0.255 (0.006)** |
| | $R_g$ | 0.412 (0.026) | 0.437 (0.017) | **0.305 (0.010)** |
|  | RMSD | 0.553 (0.044) | 0.376 (0.031) | **0.330 (0.023)** |
|  | 2D TICA | 0.603 (0.009) | 0.542 (0.002) | **0.403 (0.000)** |

#### Ablation studies on NF architecture

We conduct comprehensive ablation on the role of the latent representation, normalizing flow architectures, and coupling-mask design in Table 4. We compare four variants: TARFlow trained directly on Cartesian backbone coordinates (PROSE baseline), TARFlow trained on the learned continuous latent space, a ConformFlow variant with a four-mask coupling schedule (odd–even–first half–second half), and the final ConformFlow model with stacked odd–even affine coupling blocks. Additional ablation studies details are provided in Appendix B.3.

**Table 4.** Ablation study of coordinate representation and coupling-mask design. We compare flow modeling directly in Cartesian space (PROSE baseline), TARFlow in the learned latent space, a latent ConformFlow variant with a four-mask coupling schedule (odd–even–first half–second half), and the final ConformFlow architecture with odd–even coupling blocks.

| Dataset | Quantity | Cartesian TARFlow | Latent TARFlow | Four-mask ConformFlow | ConformFlow |
| --- | --- | --- | --- | --- | --- |
| CATH1 | End-to-end | 0.271 (0.016) | 0.159 (0.003) | 0.071 (0.004) | **0.043 (0.001)** |
| | $R_g$ | 0.250 (0.018) | 0.110 (0.006) | 0.102 (0.008) | **0.085 (0.004)** |
|  | RMSD | 0.310 (0.023) | 0.124 (0.007) | 0.077 (0.007) | **0.058 (0.003)** |
|  | 2D TICA | 0.419 (0.012) | 0.190 (0.009) | 0.196 (0.008) | **0.132 (0.003)** |
| Octapeptide | End-to-end | 0.410 (0.010) | 0.256 (0.007) | 0.189 (0.005) | **0.035 (0.000)** |
| | $R_g$ | 0.381 (0.013) | 0.232 (0.009) | 0.154 (0.008) | **0.032 (0.000)** |
|  | RMSD | 0.480 (0.010) | 0.340 (0.005) | 0.284 (0.002) | **0.033 (0.000)** |
|  | 2D TICA | 0.595 (0.011) | 0.498 (0.009) | 0.491 (0.009) | **0.329 (0.004)** |
| ATLAS | End-to-end | 0.583 (0.023) | 0.445 (0.016) | 0.433 (0.014) | **0.255 (0.006)** |
| | $R_g$ | 0.603 (0.027) | 0.436 (0.021) | 0.405 (0.026) | **0.305 (0.010)** |
|  | RMSD | 0.627 (0.050) | 0.532 (0.043) | 0.523 (0.044) | **0.330 (0.023)** |
|  | 2D TICA | 0.680 (0.013) | 0.583 (0.010) | 0.569 (0.009) | **0.403 (0.000)** |

Cartesian TARFlow performs poorly across all datasets, suggesting that direct likelihood modeling in backbone Cartesian space is challenging for simple coordinate-wise affine flows. Moving TARFlow to the learned latent space substantially improves performance on CATH1, but generalization to held-out octapeptides and ATLAS remains limited. The four-mask ConformFlow variant further improves transfer performance, while the final odd–even ConformFlow architecture achieves the best distributional agreement overall. These results show that both the learned geometry-aware latent representation and the residue-level coupling architecture are important for transferable ensemble generation.

#### Sampling efficiency

ConformFlow generates samples by one inverse step through the normalizing flow, leading to substantially lower sampling cost. As shown in Table 1, ConformFlow substantially reduces wall-clock sampling time relative to BioEmu while maintaining competitive or improved ensemble fidelity. All timing experiments generate 2,000 i.i.d. samples on a single NVIDIA L40S GPU. ConformFlow uses a batch size of 64, while BioEmu uses its default protein-length-dependent batch size. We further demonstrate the sampling efficiency gain over autoregressive flows in Appendix Table 6. ConformFlow achieves substantially faster sampling, with the speedup increasing for longer protein sequences due to its parallel RealNVP-style coupling architecture.

## 5 DISCUSSION

### Limitations

ConformFlow generates ensembles through a learned continuous latent space rather than directly in Cartesian coordinates. This improves scalability and sampling efficiency, but the final ensemble quality depends on the fidelity and coverage of the encoder–decoder. Rare conformations or proteins far outside the training distribution may therefore be limited by reconstruction accuracy. Moreover, the flow provides exact likelihoods in latent space, not exact all-atom coordinate-space likelihoods.

ConformFlow is also limited by the coverage of available equilibrium MD data. Although trained on large-scale single-structure and MD datasets, the training corpus still spans only a subset of protein sequence and conformational space. Performance may degrade for very large proteins, multidomain systems, intrinsically disordered proteins, ligand-bound states, or rare transitions that are underrepresented in current datasets. Finally, the current model generates backbone N, C*α*, and C atoms only; extending it to side-chain or all-atom ensembles is an important future direction.

### Opportunities

A key advantage of ConformFlow is the combination of fast single-pass sampling and exact latent likelihood evaluation. This allows generated conformations to be scored, filtered, or combined with physical and experimental constraints. Although this work demonstrates guidance using geometric observables, the same framework could incorporate constraints from real experiments such as SAXS, FRET, NMR, or cryo-EM. More broadly, ConformFlow provides a likelihood-based prior over protein conformations that can be combined with downstream inference objectives without retraining. Future work could also couple this equilibrium ensemble model with transition models, reweighting methods, or Markov state models to recover kinetic information.

## APPENDIX A A EXPERIMENT DETAILS

### A.1 Distributional similarity evaluation

#### End-to-end distance

The end-to-end distance (EED) measures the global extension of a protein conformation. It is defined as the Euclidean distance between the terminal backbone atoms:

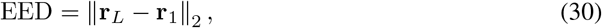

where **r**_1_ and **r**_*L*_ denote the coordinates of the first and last residues, respectively. In this work, EED is computed using *C*_*α*_ atoms.

#### Radius of gyration

The radius of gyration (*R*_*g*_) measures the overall compactness of a protein conformation. It is defined as the root-mean-square distance of atoms from their geometric center:

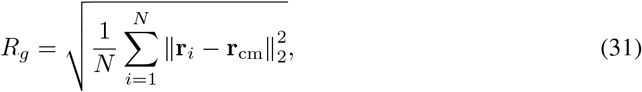

where *N* is the number of atoms, **r**_*i*_ is the coordinate of atom *i*, and **r**_cm_ is the centroid of the selected atoms. In this work, *R*_*g*_ is computed using backbone atoms.

#### Root mean square deviation

Root mean square deviation (RMSD) measures structural deviation from a reference conformation after optimal rigid-body alignment:

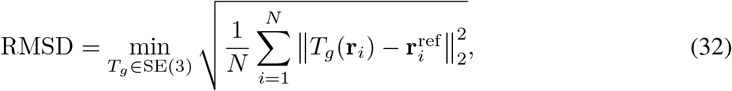

where **r**_*i*_ and 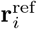 are corresponding atom coordinates in the generated and reference structures, and *T*_*g*_ is the optimal rigid-body transformation. In this work, RMSD is computed against the AlphaFold 3 predicted native structure.

#### Time-lagged independent component analysis

Time-lagged independent component analysis (TICA) extracts slow collective coordinates from MD trajectories by finding linear projections with maximal autocorrelation at lag time *τ*. It solves the generalized eigenvalue problem

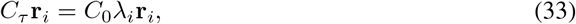

where *C*_0_ is the instantaneous covariance matrix, *C*_*τ*_ is the time-lagged covariance matrix, and *λ*_*i*_ is the autocorrelation of the *i*-th component. We fit TICA on reference MD trajectories using protein backbone torsion features and evaluate generated ensembles after projection onto the top two TICA components.

## B ADDITIONAL EXPERIMENT DETAILS

### B.1 Architecture details

ConformFlow models residue-wise continuous latents **y** ∈ℝ^*L×M*^, with *M* = 4 in all main experiments. The normalizing flow is a RealNVP-style architecture composed of 12 affine coupling blocks. Each block contains two residue-masked coupling layers: one updates odd-indexed residues while keeping even-indexed residues fixed, and the other updates even-indexed residues while keeping odd-indexed residues fixed. This odd–even masking ensures that every residue is updated once per block while preserving a block-triangular Jacobian.

The scale and translation fields in each affine coupling layer are predicted by a bidirectional masked Transformer conditioner. The conditioner receives trainable amino-acid sequence embeddings together with the frozen subset of latent variables, while the active subset is masked out to preserve exact invertibility. We use rotary positional embeddings (RoPE) over both sequence and structure-token positions to encode residue order. Each coupling network uses hidden dimension *d* = 256, 8 attention heads, and 4 Transformer layers. The Transformer blocks use pre-normalization, residual connections, multi-head self-attention, and feed-forward networks with gated nonlinear activations. The sequence embeddings, structure embeddings, Transformer conditioners, and affine output heads are trained end-to-end. The output head predicts the affine scale and shift jointly and is initialized near zero so that each coupling layer starts close to the identity map.

The full flow contains 204M trainable parameters. The model is trained by maximum likelihood on precomputed continuous latents with the structure autoencoder frozen. We optimize with AdamW using learning rate 10^*−*4^, weight decay 10^*−*4^, *β*_1_ = 0.9, and *β*_2_ = 0.95. The learning rate is linearly warmed up for 500 steps. We use gradient clipping with maximum norm 1.0 and train with bfloat16 mixed precision.

### B.2 Plug-and-play guidance details

For each protein, we draw 25 RMSD targets from the empirical MD RMSD distribution. For each target, we run 200 independent ULA chains, producing 5000 guided samples per protein. All chains are initialized from unconditional ConformFlow samples and use sampling temperature *T* = 1. We use a Gaussian constraint width of *σ* = 0.2 å . For **y**-space guidance, we run 2000 ULA steps with step size 10^*−*3^. For **z**-space guidance, we run 2000 ULA steps with step size 5 *×* 10^*−*4^. All guidance runs use 500 warm-up steps.

### B.3 Architecture ablation details

We perform an ablation study to evaluate the effect of the latent representation and coupling-mask design. Table 4 compares four variants: TARFlow trained directly on Cartesian backbone coordinates, TARFlow trained on the learned continuous latent space, a ConformFlow variant using a four-mask coupling schedule, and the final ConformFlow model with stacked odd–even affine coupling blocks.

Both TARFlow baselines use 12 affine transformation layers, matching the final ConformFlow model in depth. For Cartesian TARFlow, we remove center-of-mass translation and use random rotation augmentation during training. Our TARFlow implementation follows PROSE as closely as possible. For Cartesian coordinates, we use per-atom positional embeddings. For both TARFlow baselines, residue-level permutations are applied before each affine transformation: the first permutation in the data-to-Gaussian direction is the identity permutation, while subsequent permutations use a head-to-tail residue flip. This differs from PROSE, which uses chemistry-aware sequence permutations.

The four-mask ConformFlow variant uses six coupling blocks, with each block containing four masked coupling layers over odd, even, first-half, and second-half residue partitions. This gives the same total number of coupling layers as the final model, but changes the masking pattern. The final ConformFlow model instead uses 12 stacked odd–even affine coupling blocks, where each block alternates between odd-residue and even-residue updates.

Cartesian TARFlow performs poorly across all datasets, indicating that direct likelihood modeling in backbone Cartesian space remains difficult even after removing global translation and augmenting rotations. Moving TARFlow to the latent space substantially improves performance on CATH1, but still generalizes poorly to held-out octapeptides and ATLAS. The four-mask ConformFlow variant further improves over latent TARFlow, especially on the held-out benchmarks, but remains weaker than the final odd–even coupling design.

These results do not imply that Cartesian-space flows are inherently unsuitable for protein ensembles. Rather, they suggest that simple coordinate-wise affine transformations lack sufficient geometric inductive bias for backbone conformations. More expressive Cartesian-space flows could incorporate dense pairwise interactions or equivariant message passing, but such architectures would likely increase computational cost. Similarly, generic head-to-tail permutations and firsthalf/second-half masks provide limited protein-specific inductive bias, especially across variable sequence lengths. This supports our use of a learned geometry-aware latent representation together with odd–even residue coupling, which provides a simple, length-agnostic masking scheme while preserving efficient likelihood evaluation and sampling.

## C ADDITIONAL EXPERIMENT RESULTS

### C.1 Structure Autoencoder Evaluation

Because all downstream sampling operates in the continuous latent space **y**, the autoencoder should preserve protein backbone geometry with high fidelity. We evaluate reconstruction quality on 50 proteins from each of PDB, CATH1, CATH2, Megasim, mdcath, and ATLAS, using up to 1000 random frames per protein, and report three CA-based metrics in Table 5: RMSD, lDDT-CA (with neighbor cutoff 15 å), and CA–CA bond length (ideal 3.80 å). Across all datasets, the autoencoder achieves sub-å mean RMSD, lDDT-CA above 0.95, and CA–CA bond lengths within 0.03 å of the ideal value, indicating that both global topology and local backbone geometry are recovered with high fidelity. We therefore conclude that the learned latent representation **y** retains the backbone information needed for downstream distributional modeling.

**Table 5.** Autoencoder reconstruction quality across datasets. 50 test proteins per dataset. RMSD and lDDT measure global and local *C*_*α*_ reconstruction accuracy; the bond column reports the mean CA–CA distance (ideal 3.80 å) as a sanity check that the decoded backbone is physically valid.

| Dataset | RMSD (Å) ↓ | IDDT-CA ↑ | CA–CA bond (Å) |
| --- | --- | --- | --- |
| PDB | 0.507 (0.0155) | 0.951 (0.0040) | 3.768 (0.0080) |
| CATH1 | 0.380 (0.0071) | 0.983 (0.0001) | 3.822 (0.0002) |
| CATH2 | 0.450 (0.0118) | 0.975 (0.0002) | 3.815 (0.0002) |
| Megasim | 0.642 (0.0513) | 0.980 (0.0003) | 3.812 (0.0002) |
| mdcath | 0.508 (0.0398) | 0.974 (0.0003) | 3.797 (0.0003) |
| ATLAS | 0.763 (0.0250) | 0.960 (0.0023) | 3.797 (0.0011) |

### C.2 Fes visualization

We present additional free-energy surface (FES) visualizations on representative proteins from the CATH1 dataset. Figure 2 shows two-dimensional free-energy surfaces along the top two TICA components, while Figure 3 shows one-dimensional free-energy profiles along end-to-end distance (EED), radius of gyration (*R*_*g*_), and RMSD to the native structure.

### C.3 Parallel sampling efficiency

We further compare the scaling behavior of RealNVP-style parallel sampling against an autoregressive flow baseline. Timing is measured on a single NVIDIA L40S GPU using 32 samples per batch. As shown in Table 6, RealNVP sampling remains highly efficient as sequence length increases, while autoregressive sampling becomes progressively more expensive due to its sequential generation procedure. The speedup grows from 1.9*×* at length 8 to 38.3*×* at length 500, demonstrating the advantage of parallel invertible coupling layers for long protein sequences.

**Figure 2.**
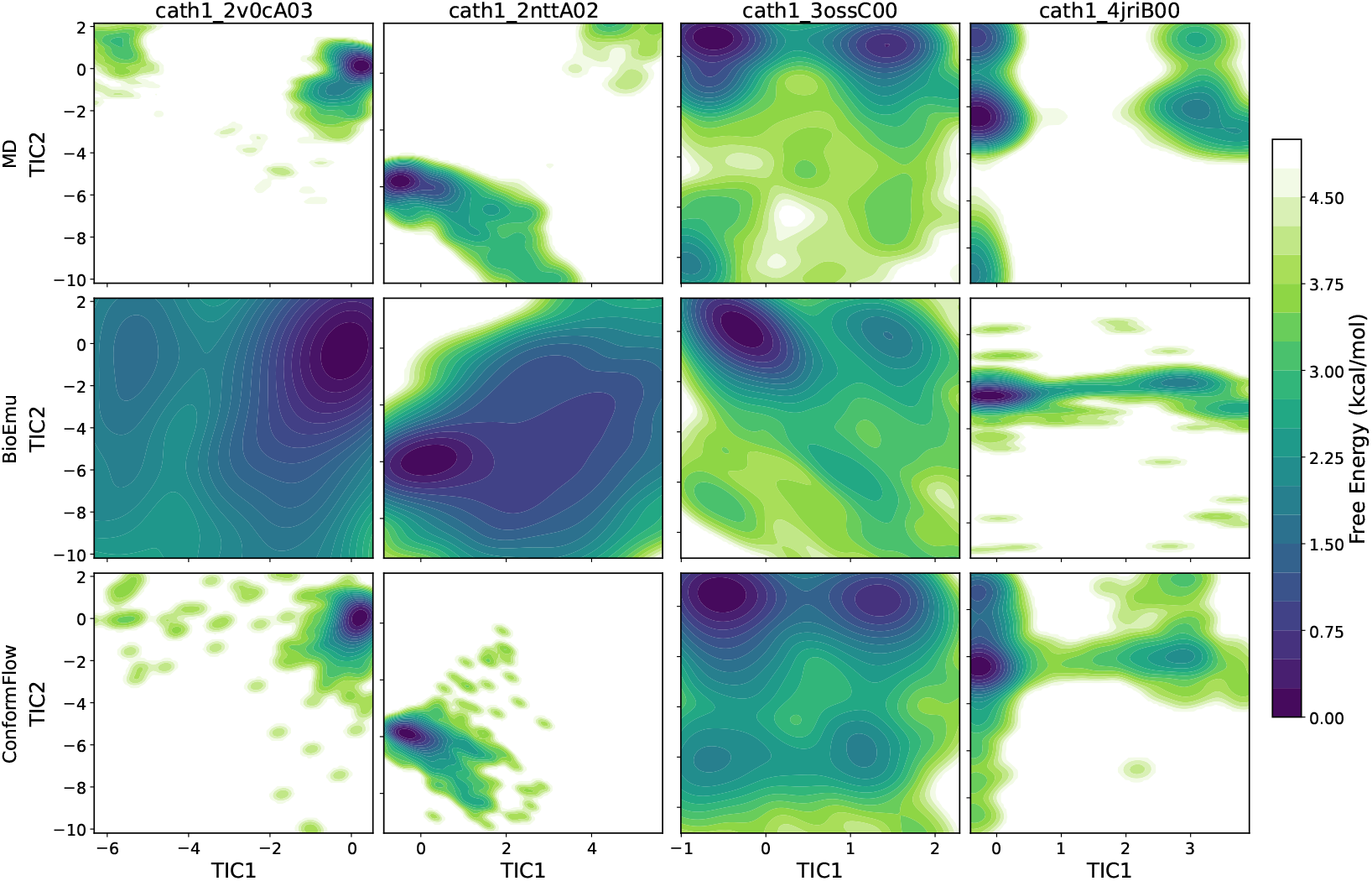
Free energy surfaces on representative CATH1 proteins along the top two TICA components. The TICA model is parameterized using *C*_*α*_ pseudo-torsion angles from the reference MD simulations, and the resulting projection is applied to conformational ensembles generated by ConformFlow and the BioEmu baseline.

**Table 6.** Sampling time comparison between RealNVP-style parallel sampling and an autoregressive flow baseline. Timing is measured on a single NVIDIA L40S GPU with 32 samples per batch. Lower time is better.

| Sequence length | RealNVP (s) | AR (s) | Speedup |
| --- | --- | --- | --- |
| 8 | 0.131 | 0.245 | 1.9× |
| 16 | 0.142 | 0.481 | 3.4× |
| 32 | 0.162 | 0.954 | 5.9× |
| 48 | 0.193 | 1.454 | 7.5× |
| 64 | 0.219 | 1.930 | 8.8× |
| 96 | 0.257 | 2.850 | 11.1× |
| 128 | 0.344 | 3.947 | 11.5× |
| 192 | 0.524 | 8.815 | 16.8× |
| 256 | 0.732 | 15.918 | 21.7× |
| 384 | 1.199 | 35.348 | 29.5× |
| 500 | 1.640 | 62.767 | 38.3× |

**Figure 3.**
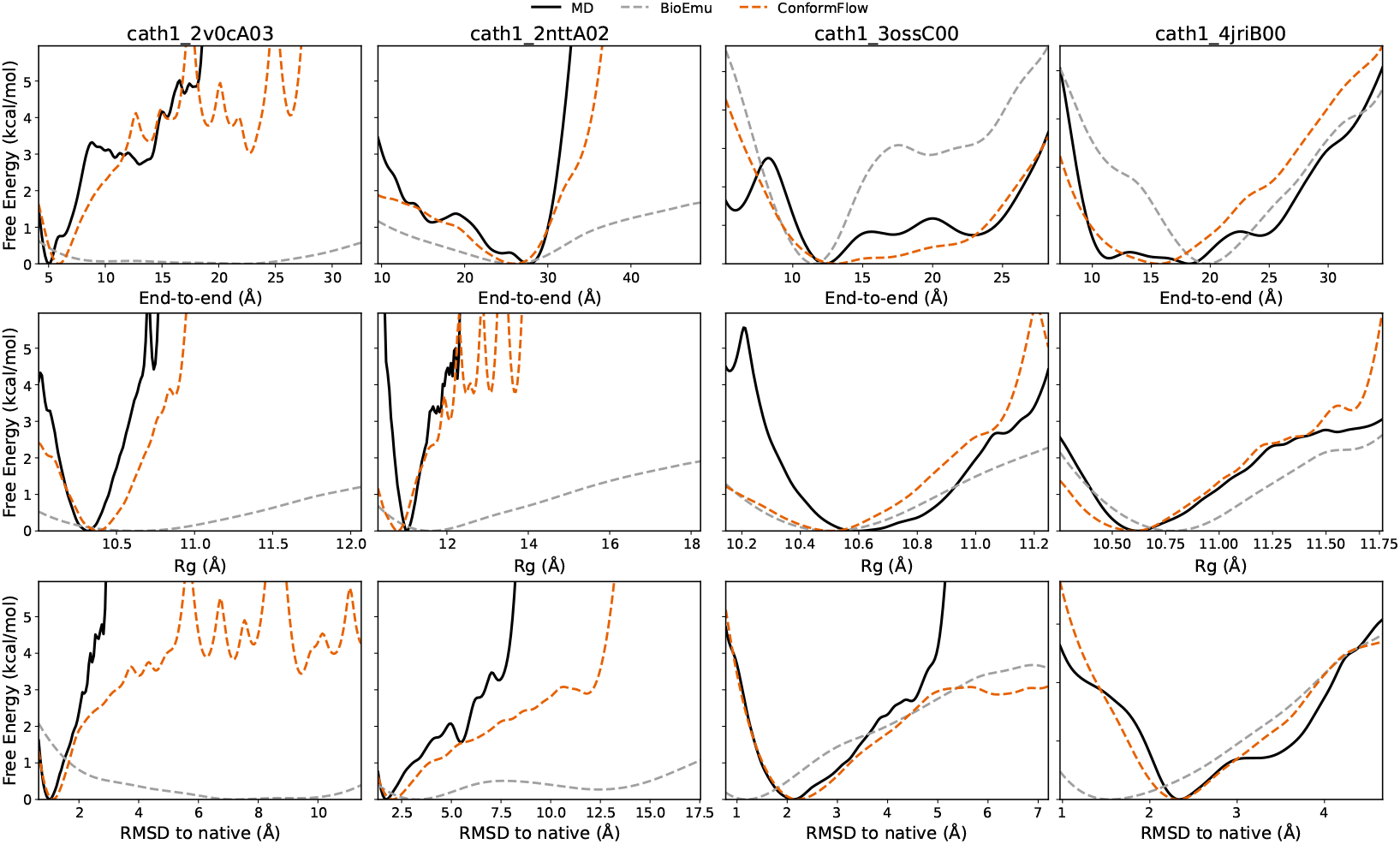
One-dimensional free energy profiles on representative CATH1 proteins along end-to-end distance, radius of gyration (*R*_*g*_), and RMSD with respect to the native structure, constructed from reference MD simulations, ConformFlow samples, and BioEmu samples.

